# Cortical networks with multiple interneuron types generate oscillatory patterns during predictive coding

**DOI:** 10.1101/2024.10.27.620494

**Authors:** Kwangjun Lee, Cyriel M.A. Pennartz, Jorge F. Mejias

## Abstract

Predictive coding (PC) proposes that our brains work as an inference machine, generating an internal model of the world and minimizing predictions errors (i.e., differences between external sensory evidence and internal prediction signals). Theoretical models of PC often rely on high-level approaches, and therefore implementations detailing which neurons or pathways are used to compute prediction errors or adapt the internal representations, as well as their level of agreement with biological circuitry, are currently missing. Here we propose a computational model of PC, which integrates a neuroanatomically informed hierarchy of cortical areas with a precise laminar organization and cell-type-specific connectivity between pyramidal, PV, SST and VIP cells. Our model efficiently performs PC, even in the presence of external and internal noise, by forming latent representations of naturalistic visual input (MNIST, fashion-MNIST and grayscale CIFAR-10) via Hebbian learning and using them to predict sensory input by minimizing prediction errors. The model assumes that both positive and negative prediction errors are computed by stereotypical pyramidal-PV-SST-VIP circuits with the same structure but different incoming input. During sensory inference, neural oscillatory activity emerges in the system due to interactions between representation and prediction error microcircuits, with optogenetics-inspired inactivation protocols revealing a differentiated role of PV, SST and VIP cell types in such dynamics. Finally, our model shows anomalous responses to deviant stimuli within series of same-image presentations, in agreement with experimental results on mismatch negativity and oddball paradigms. We argue that our model constitutes an important step to better understand the circuits mediating PC in real cortical networks.

**Author summary:** Predictive coding (PC) suggests that the brain constantly generates expectations about the world and updates these expectations based on incoming sensory input. While this theory is widely accepted, we still lack detailed models that show how specific neurons and brain circuits might carry out these processes. Here, we present a computational model which addresses this gap by including biologically plausible brain circuitry with specific types of neurons (pyramidal, PV, SST, and VIP cells) and their connections. It efficiently learns to form internal representations of visual information and uses them to predict sensory input, adjusting its predictions when errors occur. We found that particular types of neurons play different roles in these processes, and that neural oscillations emerge during the training and inference processes. Our model also replicates neural patterns observed in experiments where unexpected stimuli appear. By integrating anatomical and functional details, our work brings us closer to understanding how the brain uses predictive coding at the circuit level.

## Introduction

With only indirect access to the external world via information from the senses, the brain must infer the properties of objects based on the neural responses they elicit. The inferential nature of the perceptual experience opens the door to ill-posed problems when dealing with noisy and ambiguous sensory signals. A potential solution to the problem of maintaining a seamless perceptual experience is provided by predictive coding (PC), which proposes that the brain constantly generates and updates an internal model of the world to predict incoming sensory inputs (Mumford, 1992; Rao & Ballard, 1999; Srinivasan et al., 1982).

Central to PC is the idea that cortical circuits minimize the discrepancy between the actual and predicted sensory inputs (i.e., the prediction error). This working principle has been explored in many experimental and computational studies on PC (Attinger et al., 2017; Ayaz et al., 2019; Bastos et al., 2012; Brucklacher et al., 2023; Dora et al., 2021; Edwards et al., 2017; Eliades & Wang, 2008; Friston, 2005; Keller et al., 2012; Keller & Hahnloser, 2009; Lee et al., 2024; Ororbia & Kifer, 2022; Salvatori et al., 2021; Schultz & Dickinson, 2000; Spratling, 2010; Whittington & Bogacz, 2017; Zmarz & Keller, 2016; Lee et al., 2024; Pennartz et al., 2019). Despite early efforts to identify canonical cortical circuits for predictive coding (Bastos et al., 2012), implementations have often followed functional rather than neurobiological guidelines. Thus it remains unclear how PC may be carried out by cortical networks. Recently, progress has been made in the identification of different inhibitory interneuron types in neocortex, such as Parvalbumin- (PV), somatostatin- (SST), and/or vasoactive intestinal peptide (VIP) expressing cells, as key players in microcircuits computing prediction errors (Attinger et al., 2017; Hertag & Clopath, 2022; Hertag & Sprekeler, 2020; Hertäg et al., 2023; Keller & Hahnloser, 2009; Keller & Mrsic-Flogel, 2018). Several studies suggest that computations might happen independently for positive prediction errors (i.e. when sensory signals exceed the internal prediction) and negative prediction errors (i.e., when internal prediction exceeds sensory signals) via different circuits (Hertag & Sprekeler, 2020; Jordan & Keller, 2020; Keller et al., 2012; Keller & Mrsic-Flogel, 2018; O’Toole et al., 2023; Lee et al., 2024), providing insight in the underlying PC circuitry beyond classical frameworks. However, we are still missing a complete mapping of PC circuitry using neurobiologically plausible principles, i.e., not only the local circuitry for computing prediction errors comprising pyramidal, PV, SST, and VIP cells but also the circuitry for embedding those local configurations in realistic hierarchical cortical networks and adapting the brain’s internal world models as a result of such predictions.

In the present study, we integrate neuroanatomically informed projections patterns (Douglas & Martin, 1991; Felleman & Van Essen, 1991) with the laminar organization and cellular diversity of neural circuits in sensory cortex to provide a fully-fledged PC model for visual perception. Our model is constituted by a hierarchy of interconnected cortical areas in which positive and negative prediction errors are computed in two coexisting circuits located in superficial cortical layers of each area. Both circuits are described by the same minimal connectivity pattern (i.e., a closed pyramidal-PV loop and a VIP → SST → pyramidal disinhibitory circuit (Isaacson & Scanziani, 2011), but differ in the particular cell types targeted by incoming synapses from other areas. Upon presenting visual input to our model and subjecting it to Hebbian plasticity, we demonstrate that this architecture forms latent representations of naturalistic images (MNIST, fashion-MNIST and grayscale CIFAR-10) and reconstructs input images while minimizing prediction errors during sensory inference, and that its performance is robust against both external and internal sources of noise. The inspection of the resulting neural dynamics reveals the emergence of neural oscillations during sensory inference, with a frequency which may range from alpha (10 Hz) to gamma (>30 Hz), depending on hyper-parameter settings. These oscillations are of the stable node (or ‘damped oscillator’) type and emerge due to the recurrent interaction between representation and error neurons. The emergence of oscillations reflects the neurobiologically realism of the model and agrees with recent proposals linking alpha-band oscillations to top-down predictions (Alamia & VanRullen, 2019). When presented with a series of the same stimuli interrupted by occasional deviant stimuli, our model also displays anomalous responses to deviants as reported for standard oddball paradigms (Czigler et al., 2006; Kimura et al., 2009; Maekawa et al., 2005; Naatanen et al., 1978; Stagg et al., 2004; Stefanics et al., 2011; Tales et al., 1999; Casado-Roman et al., 2020; Parras et al., 2017), with the strength of the anomalous response depending on the position of the deviant in the series. Finally, selective inactivation of each cell type in the network (mimicking optogenetic experiments) reveals that interneurons have differential roles in PC: while PV provides a blanket of inhibition dampening network activity, SST and VIP serve to control the amplitude of neural oscillations and the number of cycles between representation and error neurons in opposing directions. Taking all these results together, the present model provides a neurobiologically plausible architecture for PC, highlighting the role of different interneuron types and the resulting oscillatory dynamics.

## Results

### Cortical circuits for predictive coding

We built a network model with circuit level detail to perform PC by incorporating canonical cortical circuits that consist of excitatory pyramidal cells and three types of inhibitory interneuron: PV, SST, and/or VIP cells (Attinger et al., 2017; Pfeffer et al., 2013) - instead of the arbitrary computational units of prediction error and representation units from previous work (Rao & Ballard, 1999).

Prediction error microcircuits computed the discrepancy between bottom-up sensory inputs and top-down predictions. These prediction errors (PEs) are positive when bottom-up sensory inputs are greater than top-down predictions and negative if vice versa (BU > TD versus BU < TD, respectively; Fig. 1A). To reflect the non-negative nature of spike signalling in the brain, PEs were further divided into positive and negative subgroups. Empirical evidence supports such grouping of neurons (Jordan & Keller, 2020; Keller & Mrsic-Flogel, 2018; O’Toole et al., 2023). Our previous work (Lee et al., 2024) also showed that implementing separate neuronal populations for positive and negative prediction errors indeed can implement PC in spiking neural networks.

**Figure 1.**
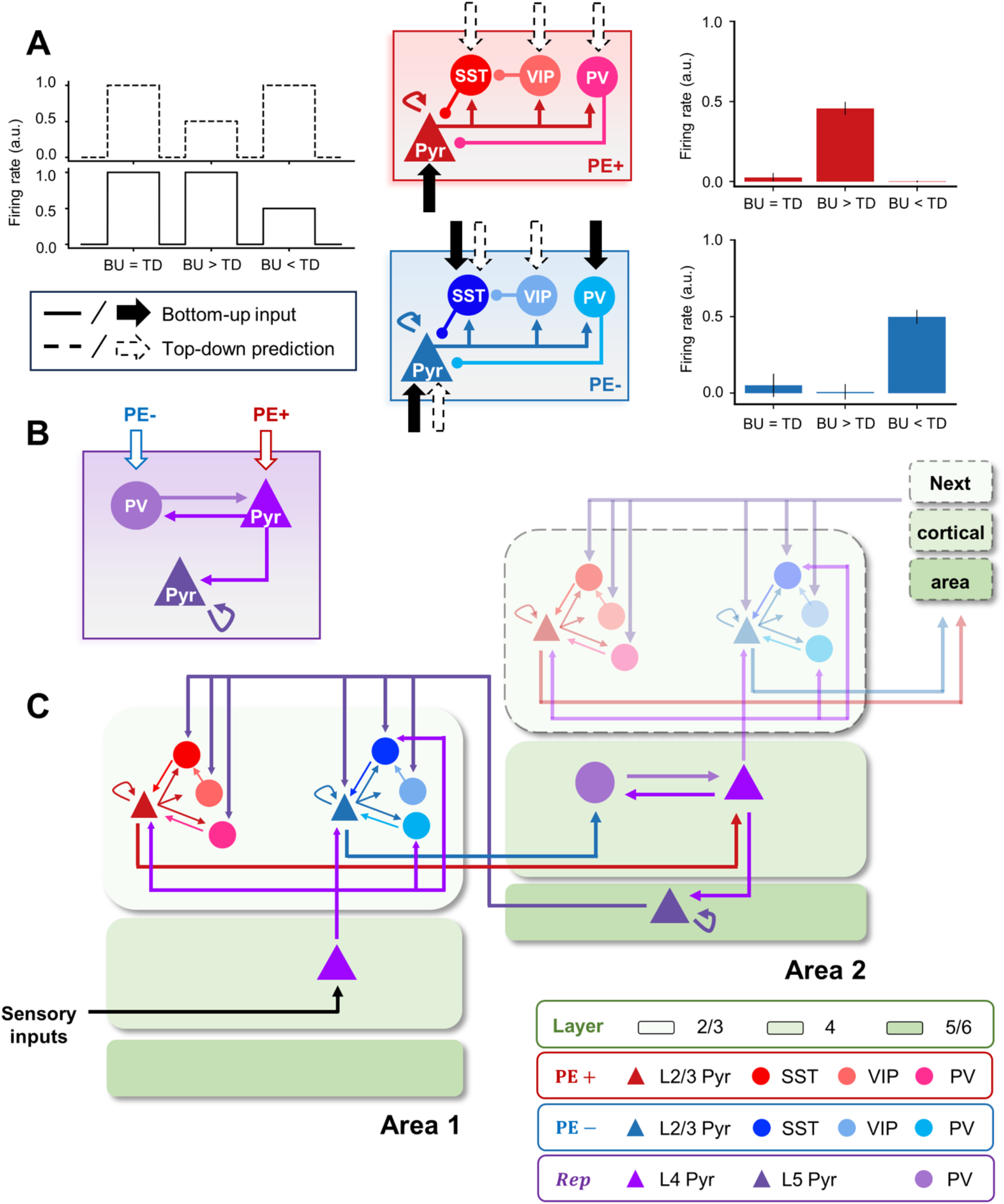
Cortical column model of predictive coding. The cortical column model of predictive coding consists of prediction error and representation microcircuits. (A) The connectivity patterns among four cortical neuron types (pyramidal, PV, SST, and VIP) within two types of prediction error microcircuits as well as synaptic inputs each of those neurons receive were determined via a combinatorial search. The canonical microcircuits (middle panel) have the same within-circuit connectivity but compute either positive (BU > TD) or negative (TD < BU) prediction errors (right panel; shown in red and blue, respectively), depending on synaptic input patterns. (B) The representation microcircuit receives the two types of prediction errors using pyramidal and PV cells. The recurrent connection between them enables the integration of the two errors to update the latent representation of sensory inputs. (C) The laminar positioning of prediction error and representation microcircuits mediating predictive coding of sensory inputs. Prediction error microcircuits are situated in layer 2/3, whereas the representation microcircuit is in layer 4/5. A higher area (e.g., Area 2) receives positive and negative prediction errors from layer 2/3 of a lower area (e.g., Area 1) to update internal representations and make predictions of incoming sensory inputs in a lower area via feedback projections from layer 5 pyramidal cells. While the model restricts the cortical hierarchy to two levels, it can be expanded to multiple columns as depicted in the upper right opaque image in Area 2 and the next cortical region.

To determine the connectivity and synaptic input patterns of prediction error microcircuits, we ran a combinatorial search to identify those circuits that allow exclusive encoding of positive or negative prediction errors. Our results showed that, despite sharing the same connectivity pattern among the four cell types within their circuits, the differences in sensory input patterns ensured that pyramidal cells from the two subgroups increased firing rates to exclusively encode either type of error (Fig. 1A).

Meanwhile, the representation microcircuit was hard-wired to continuously generate and update latent representations of bottom-up inputs (Fig. 1B). The recurrent connectivity between layer 4 pyramidal and PV cells integrated the two types of errors. Layer 5 pyramidal cells projected back top-down predictions of those bottom-up inputs to lower areas. This circuit is able to form internal representations (in the form of spatial activity patterns) of sensory stimuli and generate internal predictions to match them.

A key contribution of PV, SST and VIP interneurons is that they play an important role in generating the positive and negative prediction error responses (the rightmost panels of Fig. 1A). For example, for negative prediction error circuits, the pyramidal activity gets strongly suppressed by PV and SST activity when bottom-up input is stronger than top-down. However, if top-down input is stronger, VIP activity suppressed SST cells and the excitatory drive of pyramidal cells may overcome PV activity, resulting in a strong activity reflecting a negative prediction error. For the case of positive prediction error circuits, strong bottom-up input triggers a significant positive prediction error response (i.e., bottom-up input larger than top-down), while for strong top-down input, the activation of PV cells and the lack of any SST effects (due to VIP suppression) leads to an inhibition of pyramidal activity. In practice, the level of bottom-up and top-down signals will determine whether positive or negative prediction errors are generated, as well as their strength.

### Laminar placement of cortical circuits

Our model consisted of two adjacent cortical areas positioned along the visual cortical hierarchy (Fig. 1C). Each area in the cortical hierarchy corresponds to a cortical column that consists of supra-granular layers (i.e., layer 2/3) harboring PE microcircuits, and granular and infra-granular layers (i.e., layer 4/5) containing representation microcircuits. The laminar position and projection patterns within and between cortical columns followed the anatomy of the neocortex (Bastos et al., 2012; Douglas & Martin, 1991).

In the first, lower cortical area (e.g., V1), thalamic sensory inputs are fed to layer 4 pyramidal neurons (black triangle; Fig. 1C) and relayed to layer 2/3. Predictions regarding these bottom-up inputs are transmitted from layer 5 of the second, higher cortical area (e.g., V2; purple pyramidal cell). The disparities between the actual and predicted sensory inputs (PE+ and PE-) are computed in layer 2/3 of the first area and conveyed back to layer 4 of the second column as feedback signals (red lines) to refine latent representations of the bottom-up inputs.

Note that scaling beyond the second cortical column only requires stacking up the PE feedback loop (see the upper right part of Fig. 1C with a lower opacity). However, we limited the cortical hierarchy to two areas to emphasize the laminar architecture and cortical circuitry with interneurons that mediate predictive coding.

### Reconstruction of sensory inputs via predictive coding

To test whether our model can perform perceptual inference and learning of sensory inputs in the present architecture, which includes laminar organization, cortical circuits with interneurons, and anatomical projection patterns, we trained it with a well-known image dataset (CIFAR-10 dataset; Fig. 2A) and examined its performance in prediction error minimization (Fig. 2B) and image reconstruction (Fig. 2C).

**Figure 2.**
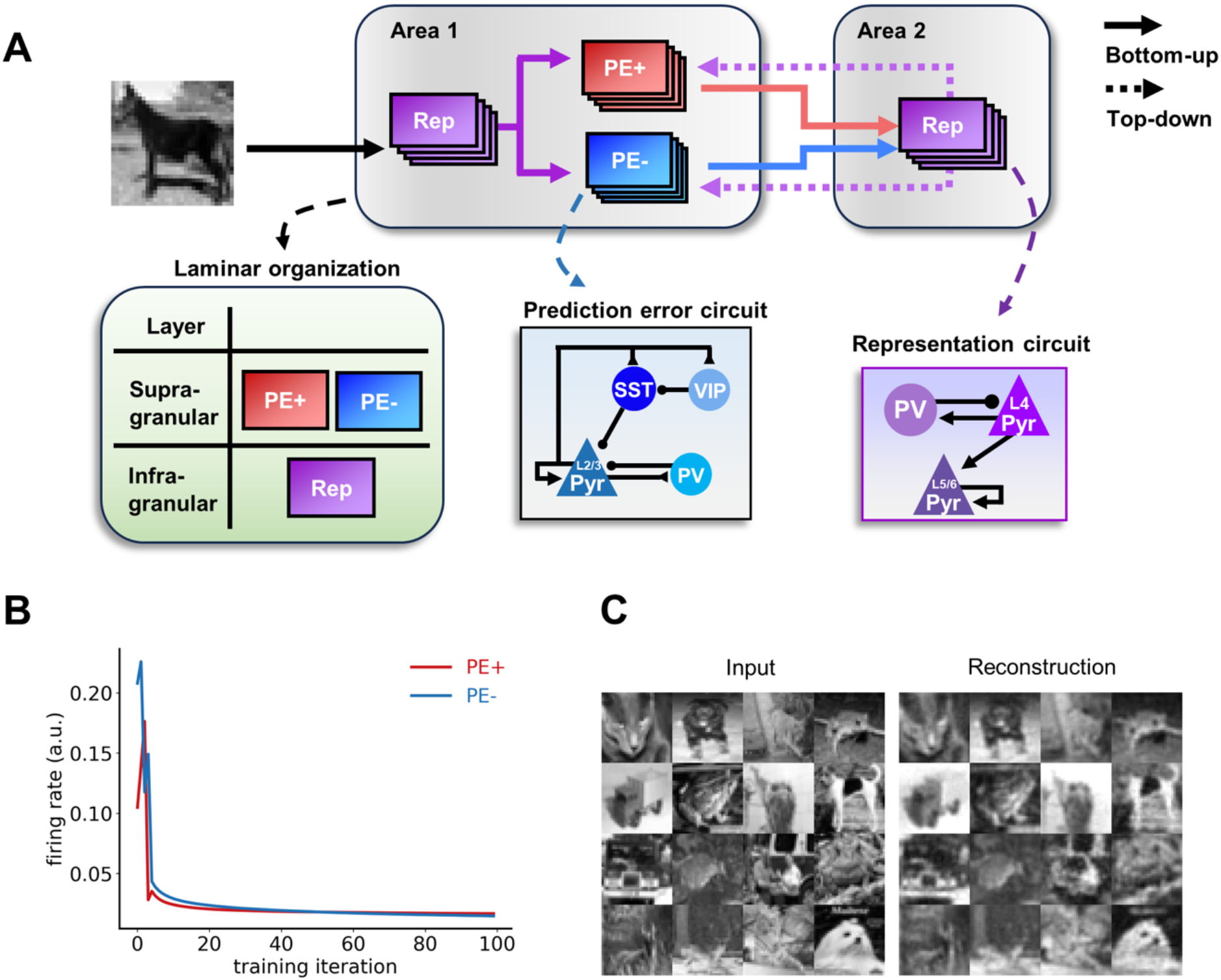
Perceptual inference and learning of naturalistic images. (A) A summary schematic of the cortical column model of predictive coding, with details on the laminar organization, prediction error circuits and representation circuits. (B) Prediction error minimization across training iterations indicates that representation learning is taking place. Both positive and negative prediction errors decrease monotonically (red and blue lines, respectively). Note that the lines indicate pyramidal cell activity from each prediction error microcircuit. (C) The model reconstructs images unseen during training, despite having been presented with only a small subset of the entire image dataset (containing 4.3% of total training images). Our model, embedded with biological constraints like laminar structure, anatomical projections and neuronal diversity, was therefore able to extract the underlying statistics from naturalistic images and learned to generate latent representations.

Except for the initial fluctuations, the monotonically decreasing prediction errors across training iterations (Fig. 2B) as well as the generative capacity to reconstruct images, not only those presented during training but also those never presented before (Fig. 2C) indicate that the model had learned the underlying statistical regularities of naturalistic images and developed latent representations thereof.

### Robustness against noise

To examine whether our architecture can acquire a robust internal model that can withstand a moderate level of sensory noise or synaptic weight jitters, we introduced Gaussian noise either to input images (i.e., external noise; Fig. 3A) or to synaptic weights within each microcircuit (i.e., internal noise; Fig. 3B).

**Figure 3.**
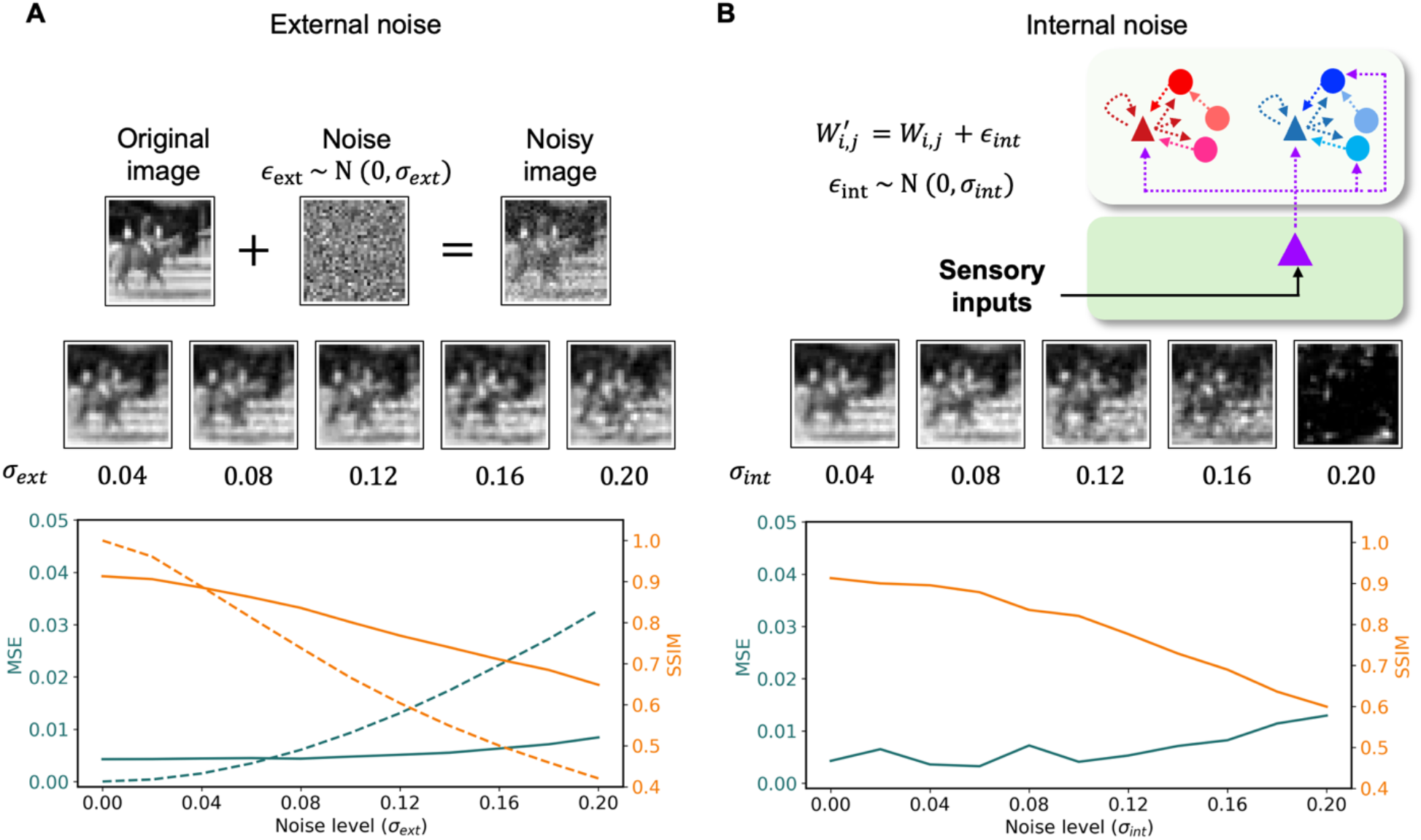
Robustness against noise. The robustness of reconstruction performance was tested against two types of noise: (A) external noise, *∈*_*ext*_ ∼ *N*(0, *σ*_*ext*_), was added to the sensory input; (B) internal noise, *∈*_*int*_ ∼ *N*(0, *σ*_*int*_), was added to synaptic weights with Gaussian noise. Despite decreases in SSIM and increases in MSE values, the model was able to infer the causes of sensory inputs up to high levels of external and internal noise. Reconstructed images with different levels of noise are shown in the middle of each panel. Orange lines indicate the structural similarity index measure (SSIM). Dark teal lines indicate the mean squared error (MSE). Solid lines compare between original images and the images reconstructed from the noisy images, whereas dotted lines compare between the original and noisy images. Note that we did not compare between the original and noisy images when adding internal noise, as it has no effect on images themselves.

When presented with images with external noise, the reconstructed images matched the original images better than the actual noisy inputs it had been provided with (solid orange line above dotted orange line, SSIMrecon > SSIMinput for noise levels > 0.05, and solid dark teal line below dotted dark teal line, MSErecon < MSEinput for noise levels > 0.06; Fig. 3A). The low MSE values and high SSIM values up to 16% of noise level to within-circuit synaptic strengths suggest that the model was also able to withstand a moderate level of synaptic jitter (Fig. 3B), which is a salient feature of biological neural networks reflecting internal synaptic heterogeneity (Song et al., 2005).

### Emergent oscillations

A close inspection of neural activity at the population level revealed that oscillatory behaviors emerge in prediction error and representation microcircuits. As prediction errors decreased (red and blue lines; Fig. 4A), stable latent representations of sensory inputs were formed (purple line; Fig. 4A) and coherent predictions of incoming sensory inputs were generated (see images on top; Fig. 4A). The rhythmic activities between the three microcircuits were phase shifted (Fig. 4B): the phase differences remained constant across time.

**Figure 4.**
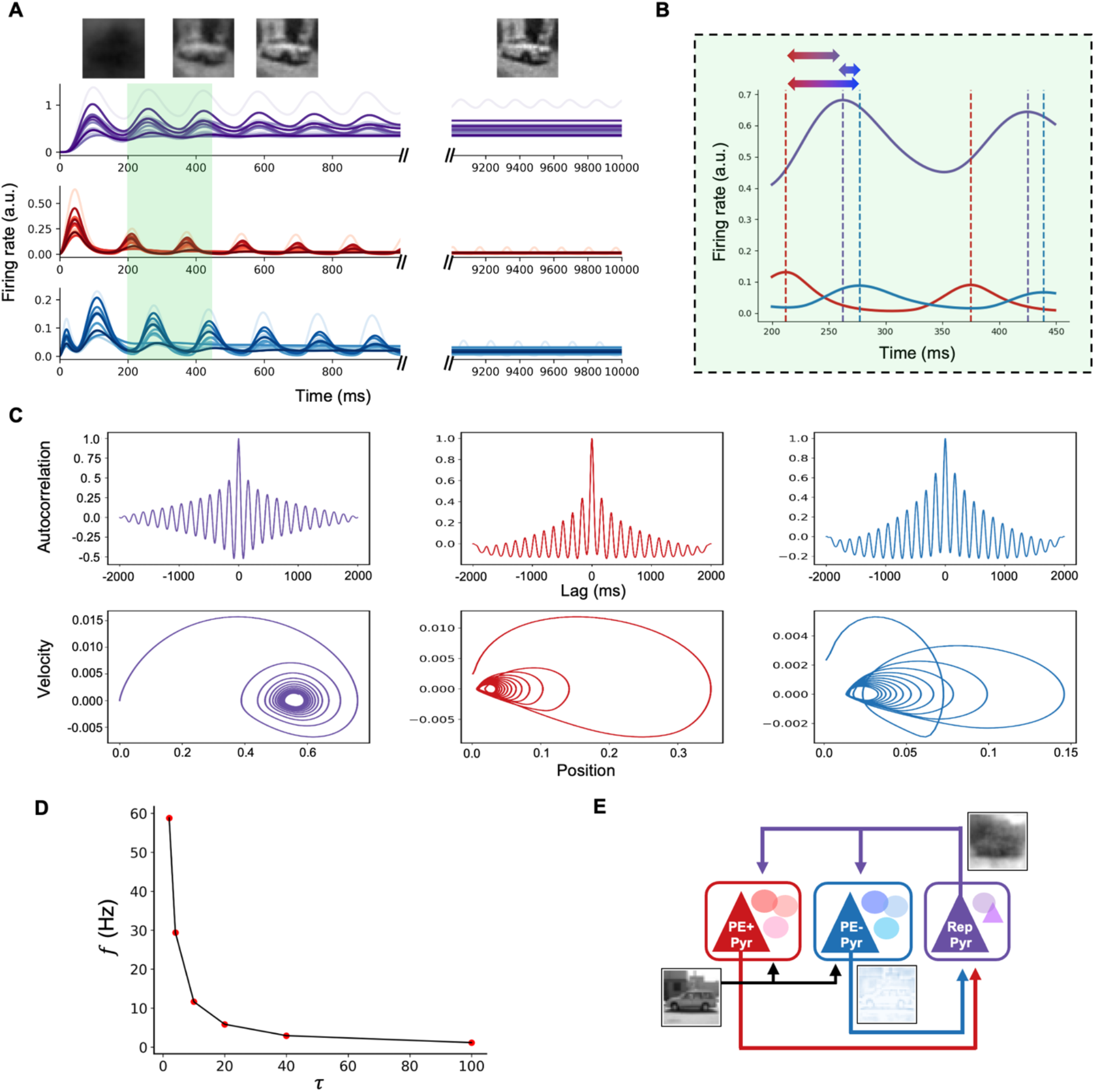
Emergent oscillations in population activity. (A) The population activity of pyramidal cells in representation (purple line; top panel), positive (red line; middle panel) and negative (blue line; bottom panel) prediction error microcircuits exhibited rhythmic activities in response to naturalistic images. As prediction errors decrease, the activity of representation microcircuits converges to generate stable images, thereby yielding better reconstructions (top images) of incoming sensory inputs. (B) The highlighted area in (A) is enlarged to show that phase differences between the two prediction error microcircuits (bottom bidirectional arrow; red to blue dotted lines), negative prediction error and representation microcircuits (middle bidirectional arrow; purple to blue dotted lines), and positive prediction error and representation microcircuits (top bidirectional arrow; red to purple dotted lines) remain constant across time. (C) Autocorrelation and phase portraits of the pyramidal cell population responses indicate damped oscillatory activity with stable fixed points. Colors as in (A). (D) The frequency of these oscillations is dependent on the membrane time constant (*τ*). (E) The oscillatory behavior reflects the inference process of predictive coding, wherein hierarchical interaction between two prediction error (PE+ and PE-; red and blue, respectively) in area 1 and representation microcircuits in area 2 (purple) minimizes the discrepancy between actual and predicted sensory signals. The grayscale image of a car (bottom left connected to black arrows) represents sensory inputs, the blue image of the car (bottom right above the blue arrow) depicts the negative prediction error signals in Area 1 (positive counterpart not shown for brevity), and the ‘blurry’ image of the car (top right next to purple arrows) depict the prediction of sensory inputs projecting from Area 2 to 1. Note that the reconstructed image is ‘blurry’ the sensory input, because we want to show prediction error signals. After a full iteration of inference steps, the image will become clearer.

The oscillating autocorrelation plots and inward spiralling phase portraits provided quantitative evidence for the presence of oscillations in all three microcircuits (Fig. 4C). The frequencies of rhythmic activities were approximately 6 Hz for representation and positive and negative prediction error microcircuits. The decreasing amplitude across lags in autocorrelation plots suggests that these oscillations were dampened across time towards stable focuses shown in phase portraits.

While the frequency of oscillations observed from microcircuits fell in alpha (Mathewson et al., 2009) and/or theta (Fiebelkorn & Kastner, 2019; Ruikes et al., 2024) range, this largely depended on the membrane time constant of neurons (Fig. 4D). The shorter the membrane time constant was, the faster the oscillation frequency became.

The rhythmic activity observed in PC microcircuits (both during training and testing phases) and their dampening oscillatory behaviors suggest that the oscillations at the population level found in our simulations are temporal signatures of the inference process of predictive coding (cf. Alamia & VanRullen, 2019). More precisely, the oscillations in our model are generated by cortico-cortical interactions between pyramidal cells of prediction error and representation microcircuits (Fig. 4E; see Fig. S2 for example), going beyond the simplified chain-like descriptions provided in previous studies (Alamia & VanRullen, 2019). All three interneurons in both PE microcircuits also showed dampening oscillatory behaviors (see Fig. S1).

### Oddball experiment

When presented with a sequence of repetitive stimuli, interrupted by a deviant stimulus (oddball paradigm; Fig. 5A), population activities of pyramidal cells in representation microcircuit and both positive and negative error microcircuits showed an oddball effect (Fig. 5B), viz. an enhanced firing rate (on top of the dampened oscillation.

**Figure 5.**
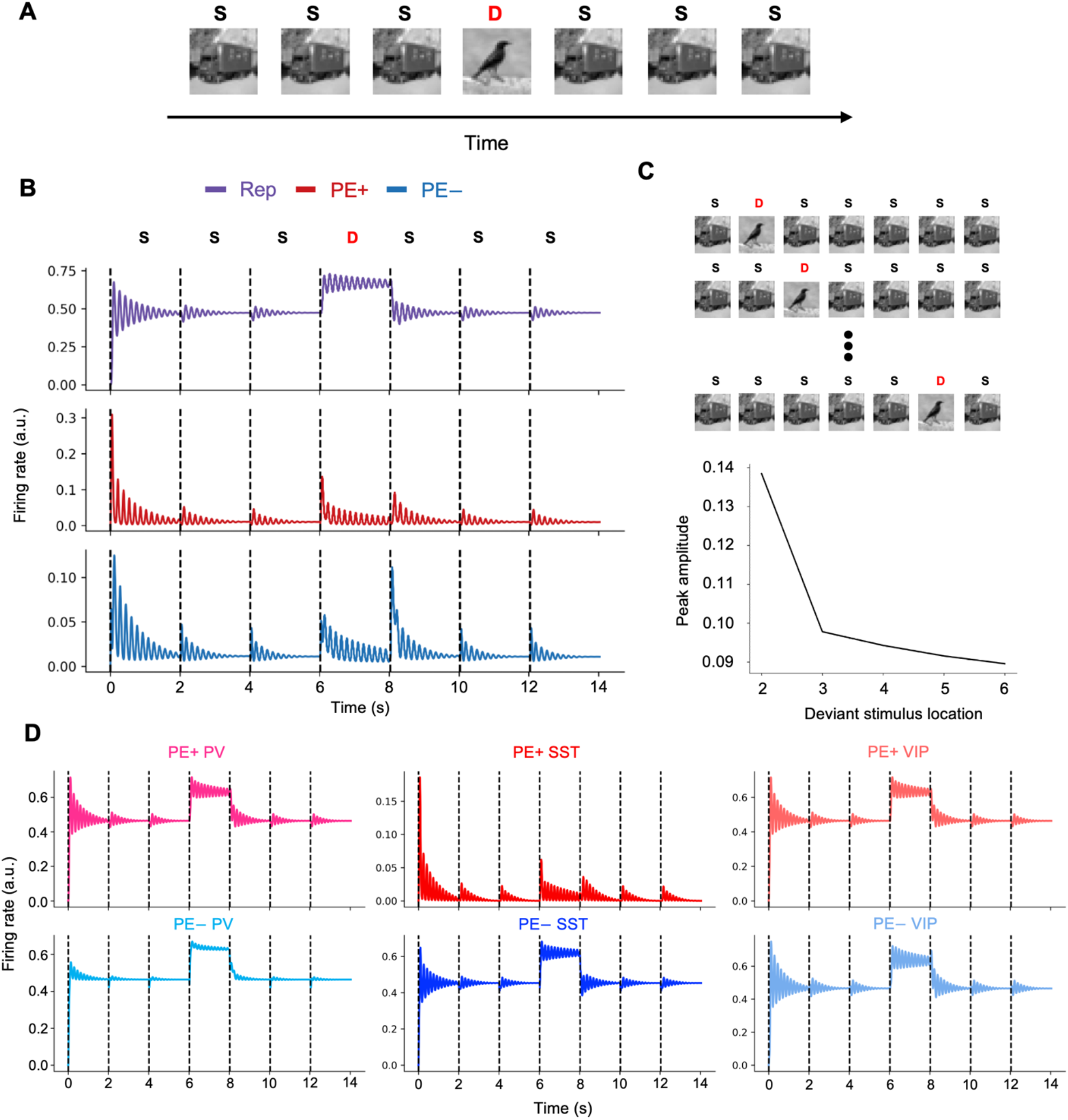
Oddball paradigm. (A) A typical oddball paradigm, in which standard stimulus, S, is repetitively presented, interrupted by a deviant stimulus, D. (B) Population activities of L5 pyramidal cells in representation (Rep; purple line) and L2/3 pyramidal cells in prediction error microcircuits (PE+ and PE-; red and blue lines) in response to the oddball sequence in (A). Prediction errors decreased over time as a consequence of the inference process. Repetitive presentation of the standard stimulus (e.g., a truck image; S) resulted in lower initial prediction errors and faster error minimization compared to the initial encounter with the stimulus (0-2 s; red and blue lines). Similarly, steady-state latent representations were reached more rapidly with repetitive presentation of a stimulus (Rep; purple line). Meanwhile, a deviant stimulus (e.g., a bird; D) elicited increased neural activity in prediction error microcircuits and led to a different steady-state representation. (C) Varying the location of deviant stimulus in the oddball sequence led to differences in initial peak amplitudes of responses in prediction error microcircuits. (D) Responses of interneurons to the oddball sequence used in (A).

Compared to the first encounter with the repetitive stimulus, the subsequent presentations generated smaller initial prediction errors, which were then minimized in a shorter amount of time (red and blue lines; Fig. 5B). As a result, latent representations converged to a stable state at an earlier time point, once the inference process during the initial encounter had been completed (purple line; Fig. 5B). The introduction of the deviant stimulus raised neural activities in prediction error microcircuits and generated higher firing rates and rhythmic activity in the representation microcircuit.

The ratio between inter-stimulus interval (ISI) and neuronal time constant (τ_exc_) had a large impact on the oddball effect: when this ratio was less than 1, the effect was more prominent than when greater than 1. The peak amplitude of prediction error activity was highest when the deviant stimulus was presented earlier in the sequence than later (Fig. 5C).

### The role of interneurons

To systematically investigate the role of interneurons in PC of sensory inputs, we set the neural activity of each of the three interneurons to zero (i.e., r_i_ = 0 where r represents firing rate of a neuron and i ∈ {PV, SST, VIP}), simulating an optogenetic silencing experiment. We analyzed population activities of the positive and negative prediction error microcircuits and the network’s performance on reconstruction of input images.

Silencing PV cells (left; Fig. 6B) resulted in a continuous increase of pyramidal cell firing rates in both the positive and negative prediction error circuits, thereby shutting down the ∼6 Hz rhythm (middle; Fig. 6B). Note that, in a more biologically realistic scenario, saturation mechanisms would place an upper bound on this firing rate increase. While inhibiting PV activities within the two PE circuits disrupted the prediction-error feedback loop between L2/3 PE and L4/5 Rep circuits, it inadvertently created a substitute loop that let the network retain the ability to predict sensory inputs. Instead of propagating positive and negative PEs, the two PE circuits simply ended up relaying bottom-up sensory inputs and top-down predictions to L4 pyramidal and PV cells of Rep circuit (black triangle and blue circle in Area 1; Fig. 1C). Integrating these activities led to a proper update of representations.

**Figure 6.**
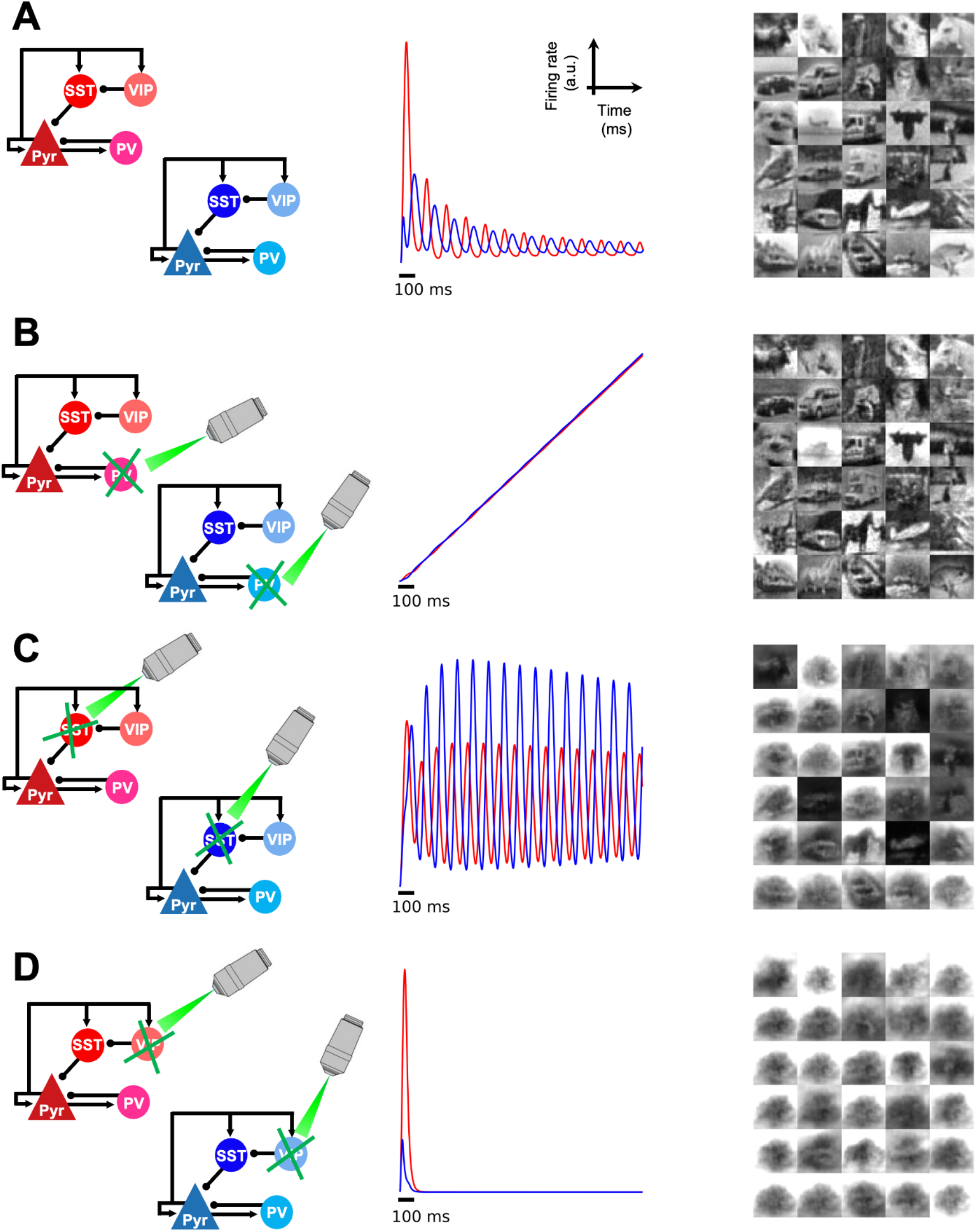
Optogenetic simulation to silence interneurons. Each of the three types of interneurons (B: PV, C: SST, and D: VIP) was silenced via optogenetic simulation to investigate their roles in predictive coding of naturalistic images. For each subpanel, the corresponding optogenetic simulation is depicted on the left, mean firing rates of pyramidal cells in positive and negative prediction error (PE) microcircuits (red and blue, respectively; in arbitrary units) is shown in the middle, and reconstructed images (i.e., predictions of sensory inputs) are shown on the right. (A) A canonical microcircuit without deactivating any interneuron is presented as a reference for comparison. Positive and negative PE responses decrease across time as a result of perceptual inference. The rhythmic activities of the two PE microcircuits were phase-shifted (see Fig 4B). (B) When PV cells were silenced, both positive and negative PE microcircuits lacked inhibitory inputs, leading to a constant increase in prediction errors. However, representation microcircuits were still able to make predictions about incoming sensory inputs, thanks to the inadvertent substitute loop that still retains the capacity to predict sensory inputs, in which the positive PE microcircuit relays bottom-up sensory inputs and the negative PE microcircuit transmits top-down predictions. The representation microcircuit integrates these two signals to update internal representations of sensory inputs. (C) Without SST cells, both positive and negative prediction error microcircuits showed persistent rhythmic activities throughout the duration of stimulus presentation, resulting in a failure to minimize prediction errors and predict incoming sensory inputs. (D) Knocking out VIP cells silenced both types of prediction error microcircuits and led to a failure to reconstruct images.

Despite silencing SST cells (left; Fig. 6C), oscillatory activities were still observed in both the positive and negative prediction error microcircuits (middle; Fig. 6C). However, these oscillations were not dampened to a fixed point. This indicates a failure to correctly infer the causes of sensory inputs, as evidenced by the model’s inability to reconstruct input images (right; Fig. 6C).

In the absence of VIP cell contributions to prediction error computation (left; Fig. 6D), pyramidal cell activities in both the positive and negative prediction error microcircuits quickly shut down (middle; Fig. 6D). Removing the VIP-SST disinhibitory circuit shifted the excitatory-inhibitory balance in synaptic inputs towards inhibition of pyramidal cells, resulting in a cessation of firing activity. Without the feedforward prediction errors that guide the proper inference of incoming sensory inputs, the network was unable to reconstruct input images (right; Fig. 6D).

## Discussion

In this work, we have demonstrated that PC may be explained in the context of a biologically realistic set of principles of cortical architectures. We have attempted to minimize the number of functional assumptions and instead root our model on neurobiologically-compatible principles (Fig. 1): hierarchically-organized relationships between cortical areas, in terms of laminar projection patterns (Felleman & Van Essen, 1991), laminar organization of microcircuits (Bastos et al., 2012; Douglas & Martin, 1991; Pennartz et al., 2019), Dale’s principle for excitatory and inhibitory cells (Dale, 1935), diversity of cell types (Keller & Mrsic-Flogel, 2018; O’Toole et al., 2023; Tremblay et al., 2016) and Hebbian learning rules (Hebb, 1949). Likewise, our model is able to perform PC efficiently (Fig. 2), but the results go beyond simply inferring the sensory input towards more neurobiologically relevant directions: good performance in the presence of external and internal noise (Fig. 3), division between positive and negative prediction error circuits (Jordan & Keller, 2020; Keller et al., 2012; O’Toole et al., 2023; Lee et al., 2024), emergence of neural oscillations during visual processing (Fig. 4) (Alamia & VanRullen, 2019; Gray & Singer, 1989; Henrie & Shapley, 2005; Perrenoud et al., 2016; Ray & Maunsell, 2011), and enhanced neural responses to deviant stimuli in mismatch negativity and oddball paradigms (Fig. 5) (Czigler et al., 2006; Kimura et al., 2009; Maekawa et al., 2005; Naatanen et al., 1978; Stagg et al., 2004; Stefanics et al., 2011; Tales et al., 1999; Casado-Roman et al., 2020; Parras et al., 2017; see Lieder et al., 2013 and Friston, 2005 for computational model and theory).

The consideration of multiple types of inhibitory neurons constitutes a fundamental ingredient in our study, and it follows previous work on the role of SST and VIP cells in generating prediction errors (Hertag & Sprekeler, 2020; Keller & Mrsic-Flogel, 2018). A large number of studies in recent years have highlighted SST and VIP cells as key players in integration of bottom-up and top-down input (Beerendonk et al., 2024; Garcia Del Molino et al., 2017; Isaacson & Scanziani, 2011; Litwin-Kumar et al., 2016; Moreni et al., 2023, 2024), thus their involvement in PC should be seriously considered. Connections between pyramidal, PV, SST and VIP cells were chosen following canonical circuit configurations common in the literature (Beerendonk et al., 2024; Garcia Del Molino et al., 2017; Litwin-Kumar et al., 2016), which include important connectivity patterns to reflect the role of inhibitory neurons in providing blanked inhibition and also disinhibitory pathways. Patterns of external projections to those neurons were, on the other hand, determined via a combinatorial search. In such a search, we indeed found different connectivity patterns which would give rise to the same behavior, which suggests some level of degeneracy –allowing for multiple connectivity combinations to yield the same performance when computing prediction errors. This level of degeneracy could be actually much higher if we relax our constraints on the microcircuit connectivity, allowing for example weaker connections to pyramidal cells which could be compensated with stronger external projections. This suggests that cortical circuits with the same functionality could indeed be formed via different mechanisms and display a diversity in circuit motifs (Hertag & Sprekeler, 2020).

Contrary to the diversity of cell types, neural oscillations were not specifically ingrained in our architecture, but rather constitute an emergent phenomenon resulting from neural dynamics due to excitatory-inhibitory interactions within microcircuits. The nature of these oscillations, of the stable-focus type or ‘damped oscillator’ type, does not constitute a problem by itself. While these oscillations will decay and disappear in an idealized system, the presence of external or internal noise is enough to maintain such oscillations in a noise-driven state and propagate them to other areas within the cortical hierarchy, as shown elsewhere (Klaver et al., 2023; Lindeman et al., 2021; Mejias et al., 2016). Indeed, the performance of our model is well preserved under low or moderate levels of external or internal noise (Fig. 3), so noise-driven oscillations are likely to survive in real conditions. Furthermore, fast oscillations such as gamma, beta, and perhaps even high alpha might be better described by noise-driven oscillations (i.e., stable focus under noise) rather than by a sinusoidal-like limit cycle, given the low amplitude and weak coherence observed in LFP recordings for these ranges (Bastos et al., 2015; Bosman et al., 2012; Henrie & Shapley, 2005). As reflected by our simulations, different interneuron types will likely play complementary roles in these oscillatory dynamics, which makes the question experimentally accessible via optogenetic inactivation protocols. While the precise frequency of the oscillations in our model depends on parameters such as population time constants, whose values are not directly accessible from recordings, the potential range includes frequencies from gamma to alpha. The latter case would be in line with previous computational work which has linked the presence of top-down signals with the appearance of alpha oscillations (Mejias et al., 2016) and particularly in the context of predictive coding (Alamia & VanRullen, 2019). What the exact role of these oscillations is within PC is a question that needs further study, although our results already suggest that they might be reflecting, or mediating, the communication between representation and error neurons during the inference process. This hypothesis is at least consistent with experimental observations of alpha and theta waves mediating inter-regional communication (Gallina et al., 2024; Zhang et al., 2018). Alpha waves, for example, are known to propagate in the feedback direction in the absence of stimuli (Bastos et al., 2015; Halgren et al., 2019; Saalmann et al., 2012; van Kerkoerle et al., 2014; Zhang et al., 2018) which could very well involve the interaction between error and representation units in our model. Theta oscillations might play a similar role, although this is less clear given that their propagation has often been linked to feedforward traveling waves in the brain (van Kerkoerle et al., 2014; Zhang et al., 2018). Meanwhile, Vinck et al. (2024) limits the role of oscillatory dynamics in predictive processing to stabilizing neural representations and facilitating plasticity whilst arguing that aperiodic transients are responsible for sensory inference.

Our model can, and should, be extended in different directions in the future. First, while we have committed to keep our working assumptions close to neurobiology, there are several aspects that could still use refinement. For example, neural dynamics has been described in terms of mass models of population activity, but spiking network models for PC can be used to further enhance realism (Lee et al., 2024). In addition, cortical microcircuits with pyramidal, PV, SST and VIP cells are idealizations, and deep layers have been modelled with less detail than superficial ones. These two aspects could be solved or alleviated by heading towards data-constrained models of cortical columns, for which several models with multiple interneurons are now available (Billeh et al., 2020; Jiang et al., 2024; Moreni et al., 2023, 2024). Considering other learning rules on top of these circuits, such as inhibitory plasticity rules, could contribute to balance excitation and inhibition and better refine prediction error circuits (Hertag & Sprekeler, 2020). Our model also assumes that the learned feedback weights (the purple arrows from L5 pyramidal cell in Area 2 to L2/3 PE circuits in Area 1; Fig. 1C) targeting positive and negative prediction error circuits are approximately the same, even though they target different neurons. While this condition may be in practice relaxed in the model and good results are obtained when these weights are moderately different, it is necessary to explore which concrete biophysical mechanism might be able to provide this soft homeostatic effect. As positive and negative prediction error circuits might coexist within the same space (O’Toole et al., 2023), local neuromodulation stands as a plausible element here. Finally, beyond adding more neurobiological features to the model, another open avenue is to expand the functionality of the model to implement more sophisticated types of task, including object invariance properties to take into account visual rotations or transformation common in real-world applications (Brucklacher et al., 2023).

Overall, the present work illustrates that PC does not require a hypothesis-driven implementation (such as a free energy minimization principle), but rather that it can be obtained by considering realistic cortical architecture, cell variability and Hebbian learning rules. According to our results, PC can emerge naturally in realistic cortical networks without additional assumptions. Our finding that oscillations emerge spontaneously in our model demonstrates a link between PC and brain oscillations tied to cortical bottom-up and top-down processing, underscoring its neurobiological plausibility.

## Methods

### Neuron model

Neurons follows a linear firing rate model (Wilson & Cowan, 1972):

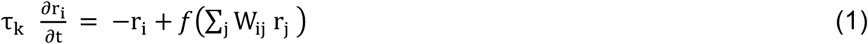

The firing rate of a neuron *i* (r_i_, where i ∈ {pyr, PV, SST, VIP}) changes according to the membrane time constant (τ_k_, where k ∈ {exc, inh}), the leak term (−r_i_), and the integration of incoming synaptic inputs (f(∑_i_ W_ij_ r_j_)) (Eq. 1). The membrane time constants for excitatory and inhibitory neurons were kept the same (20 ms) for simplicity. The activation function (*f*) was rectified linear unit (ReLU).

### Microcircuit configurations

The connectivity pattern among the four cortical cell types in positive and negative prediction error microcircuits and the synaptic input patterns to those neurons was determined via an exhaustive combinatorial search. Specifically, we looked for a canonical pattern of connectivity that exclusively computes either positive or negative prediction errors depending on the synaptic input pattern (Fig. 1A). To narrow down the search space, we imposed the following constraints based on empirical findings (Bastos et al., 2012; Pfeffer et al., 2013; Pennartz et al., 2019): 1) pyramidal cells must receive bottom-up sensory inputs (e.g., from thalamus to layer 4 in V1); 2) VIP cells must receive top-down inputs; 3) SST and PV cells both contact pyramidal cells with inhibitory synapses; 4) VIP cells play a disinhibitory role by inhibiting SST cells; and 5) pyramidal cells project to all neuron types within a microcircuit, including themselves. The strength of each synapse within all microcircuits (i.e., PE+, PE-, and Rep) was kept at one for simplicity.

From all the potential microcircuits satisfying the criteria above, we selected a biologically plausible example without loss generality from a computational perspective (Fig. 1A). Note that, however, substituting it with other set of patterns would lead to similar results.

### Learning naturalistic images

Only a small subset of the CIFAR-10 dataset (nimage = 2560, nclass = 10, and nsample = 256; 5.1% of the total images in the dataset) was used for the training. Without loss of generality, with respect to predictive coding of sensory signals, the colored images of the CIFAR-10 dataset was converted to grayscaled images to reduce the computational costs.

Using a batch training method (batch size = 64), each image was presented for 2 seconds, during which the model formed latent representations of incoming sensory inputs. Before presenting the next batch of stimuli, the network was shown an empty screen for 0.2 seconds. During this inter-stimulus interval, neural activities across the cortical hierarchy quickly reset. The model was shown each image 100 times.

The updating of synaptic weights between prediction error microcircuits and representation microcircuit occurred immediately after the stimulus presentation had ended and followed a Hebbian learning rule as in Lee et al. (2024):

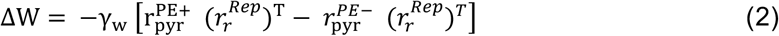

The first term within the squared bracket represents the contribution of positive errors (PE+), whereas the second term corresponds to the contribution of negative errors (PE-; Eq. 2). Note that *γ*_*w*_ represents a learning rate and 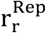 the firing rate of L5 pyramidal cells in representation circuit.

The extrinsic weights between PE and representation microcircuits, W, were symmetric and shared between the positive and negative PE microcircuits. The synapses within all microcircuits had a fixed weight of one and were not subject to synaptic plasticity.

## Conclusion

We propose a novel cortical network model with laminar and cell-type specific architecture that can perform perceptual inference learning of sensory input. We observed rhythmic activity spontaneously emerging during propagation of predictions and errors and dissected the role of different cortical neuron types in generating prediction errors.

## Acknowledgments

This work was done with the support of EBRAINS and HBP computing services. We thank Maria Nefeli Panagiotou for early iterations of this work.

## Author contributions

KL, CP, JFM conceived and designed the study; KL performed the research; KL and JFM analyzed the results; KL, CP and JFM wrote the manuscript.

## Data and materials availability

All information needed to reproduce the results of this manuscript are in the main text and Materials and Methods section, and the Python code developed will be made available upon publication of this work.

---

**Figure S1.**
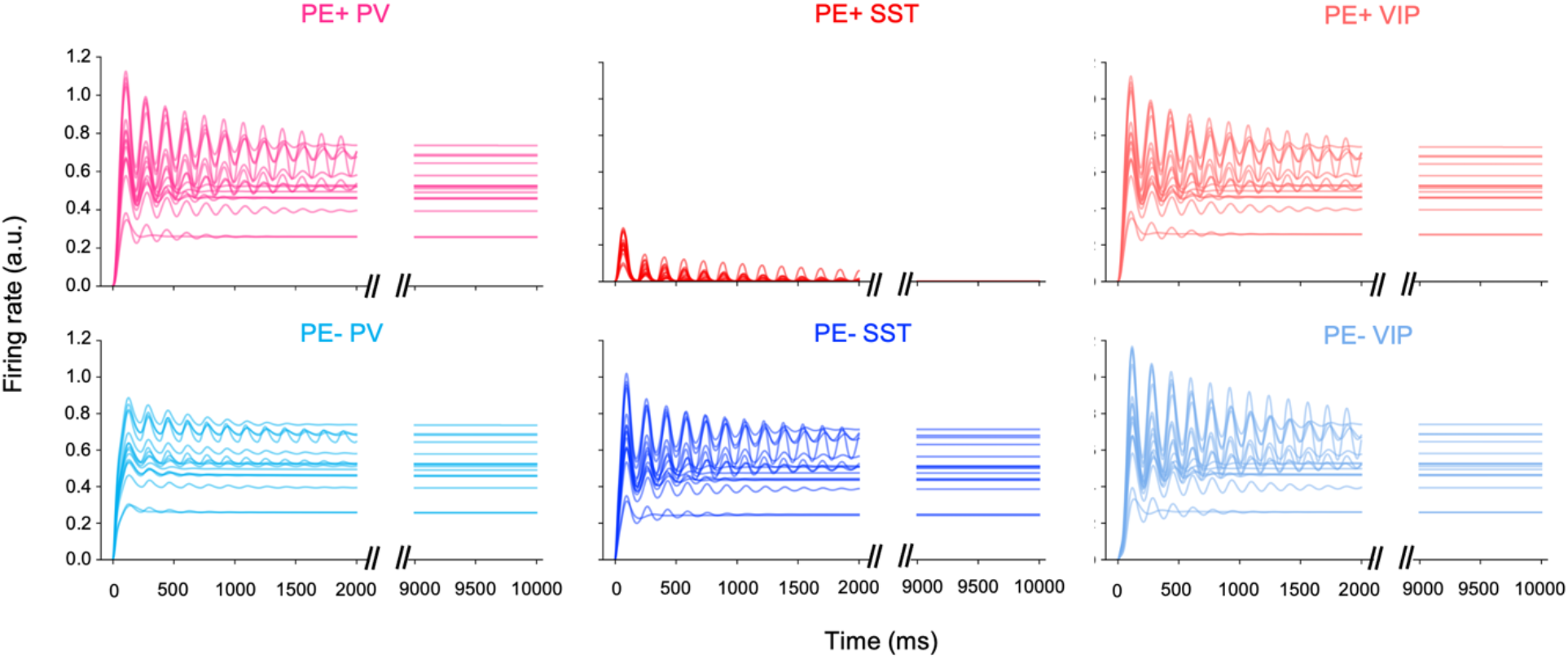
Oscillatory behavior of interneurons in response to naturalistic images. Similar to the pyramidal cells (see Fig. 4), each of the three interneurons (PV, SST, and VIP) in positive and negative PE microcircuits (color as in Fig. 1C) also show rhythmic activities that dampen across time to reach stable focuses.

**Figure S2.**
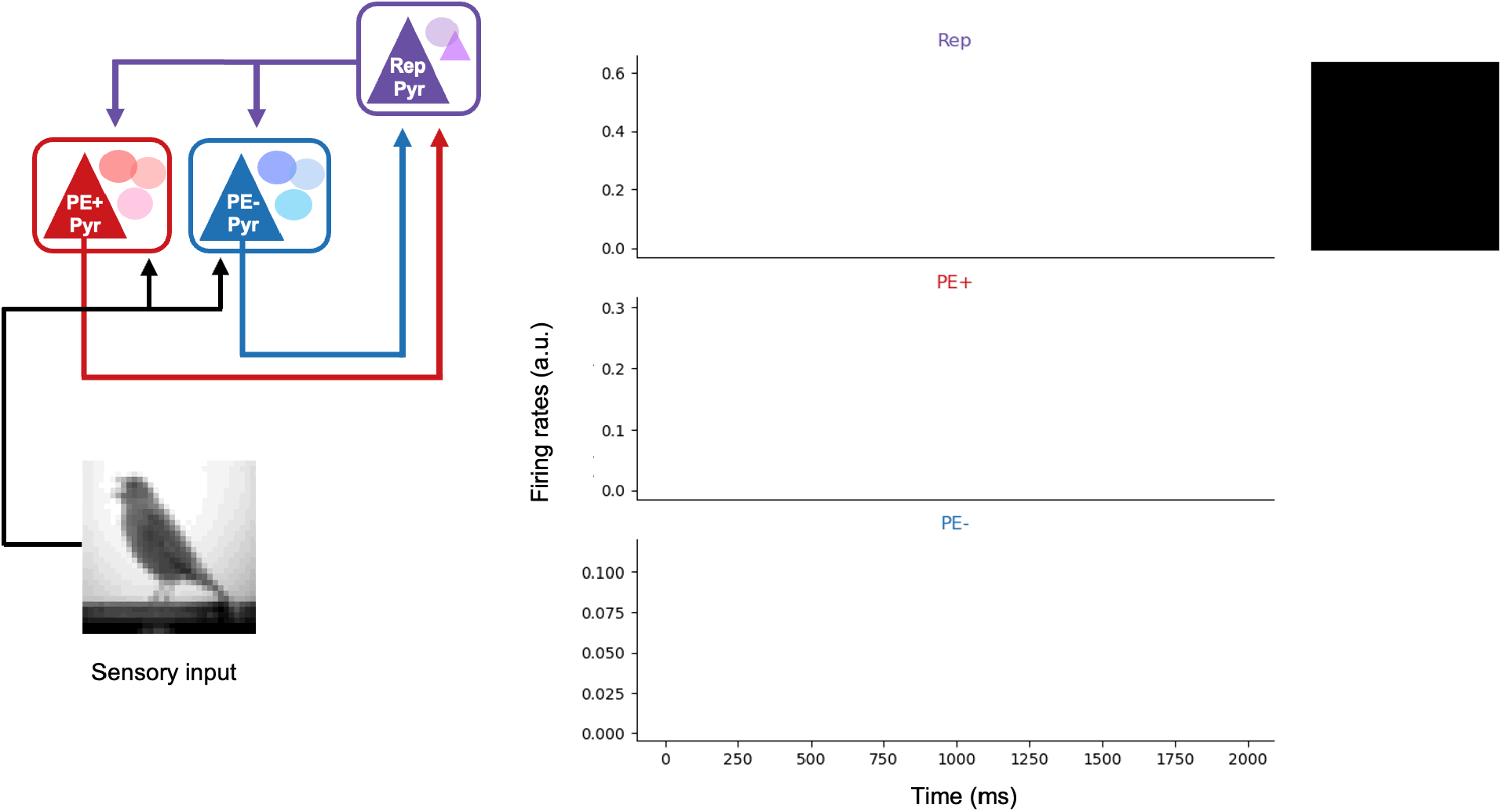
Predictive coding of sensory inputs and emergent oscillations. **(Left)** Given sensory inputs (e.g., a bird), positive and negative prediction error (PE) and representation (Rep) microcircuits hierarchically interact to infer the visual object in pursuit from sensory signals (i.e., inference). **(Right)** The three lines represent mean firing rates of pyramidal cells in each of the three microcircuits: L5 for Rep in Area 2 and L2/3 for PE+ and PE-from Area 1 (see Fig 1 for detailed architecture). For visual aid purpose, we show top-down predictions (gray animation) and bottom-up positive and negative PEs (red and blue animation, respectively) in the same dimension as original sensory inputs. As PE decreases, the color of animation becomes lighter and prediction becomes clearer. All three microcircuits show dampening oscillatory behaviors.

